# Inclusion bodies formed by polyglutamine and poly(glycine-alanine) are enriched with distinct proteomes but converge in proteins that are risk factors for disease and involved in protein degradation

**DOI:** 10.1101/2020.05.04.076547

**Authors:** Mona Radwan, Jordan D. Lilley, Ching-Seng Ang, Gavin E. Reid, Danny M. Hatters

**Author notes:** Correspondence (D.M. Hatters). Twitter: @DannyHatters.

## Abstract

Poly(glycine-alanine) (polyGA) is one of the dipolypeptides expressed in Motor Neuron Disease caused by C9ORF72 mutations and accumulates as inclusion bodies in the brain of patients. Superficially these inclusions are similar to those formed by polyglutamine (polyQ) in Huntington’s disease and both have been reported to form an amyloid-like structure suggesting they might aggregate via similar mechanisms to confer cellular dysfunction similarly. Here we investigated which endogenous proteins were enriched in these inclusions and whether aggregation-prone lengths of polyQ (Q_97_), in context of Huntingtin exon 1, shared similar patterns to aggregation-prone lengths of polyGA (101_GA_). When co-expressed in the same cell, polyGA_101_ and HttQ_97_ inclusions adopted distinct phases with no overlap suggesting different endogenous proteins would be enriched. Proteomic analyses indeed yielded distinct sets of endogenous proteins recruited into the inclusion types. The proteosome, microtubules, TriC chaperones, and translational machinery were enriched in polyGA aggregates, whereas Dnaj chaperones, nuclear envelope and RNA splicing proteins were enriched in polyQ aggregates. Both structures revealed a synergy of degradation machinery including proteins in the polyQ aggregates that are risk factors for other neurodegenerative diseases involving protein aggregation when mutated, which suggests a convergence point in the pathomechanisms of these diseases.

## INTRODUCTION

The formation of protein inclusions is a hallmark of many neurodegenerative diseases. In Huntington’s disease, amino-terminal fragments of mutant Huntingtin (Htt) protein aggregate into intraneuronal inclusions [1]. The aggregation of mutant Htt is triggered by an abnormally expanded polyglutamine (polyQ) sequence encoded in exon 1 that arises by CAG trinucleotide repeat expansions [2, 3]. Long polyglutamine sequences form cytoplasmic or nuclear inclusions in animal and mouse models and is associated with a pathological cascade of events (reviewed in [4]).

In motor neuron disease caused by *C9ORF72* GGGGCC hexanucleotide repeat expansion mutations, protein inclusions arise from the aggregation of dipolydipeptides (DPRs) expressed abnormally from the expanded GGGGCC hexanucleotide repeat sequence. 5 different DPRs are expressed, namely dipeptide polymers of glycine-alanine (polyGA), proline-arginine (polyPR), glycine-arginine (polyGR), proline-alanine (polyPA), and proline-glycine (PolyPG). Of these polyPR and polyGR are profoundly toxic when expressed in cell culture and animal models targeting mechanisms in ribosome biogenesis, translation, actin cytoskeleton among others [5-12]. PolyGA appears less toxic than the others although has been reported in some models to confer toxicity [13-21]. We previously reported polyGA to be mildly toxic to cultured Neuro2a cells and to induce a distinct network of proteome changes that occur compared to the arg-rich DPRs [12]. We also noted a distinction to the other DPRs in forming large inclusions that morphologically are similar to the inclusions formed by polyQ. Furthermore, it has been reported that polyGA inclusions are, like polyQ, SDS-insoluble and amyloid-like [22, 23]. PolyGA is also more widespread in MND-patient brain tissue compared to the other DPRs [24].

Here we investigated the proteinaceous composition of polyGA inclusions and compared the profile quantitatively to inclusions of polyQ using a Huntingtin exon 1 model (Httex1Q_97_) in a mouse neuroblastoma cell culture model using a novel proteomics-based approach to enrich the inclusions from cells under mild lysis conditions. We find distinct recruitment patterns to each inclusion type and also some similarities in biological mechanisms pertaining to degradation and a convergent pathomechanism for neurodegenerative diseases involving inappropriate protein aggregation.

## METHODS

### Plasmids

A pEGFP-based construct expressing polyGA dipeptide repeat length of 101 dipeptides (polyGA_101_) was generated as described previously [12]. This construct expresses a GFP fusion tag at N-terminus of the polyGA. pT-REx vector expressing exon 1 of Htt (Httex1) with polyQ sequence length of 97 and C-terminal mCherry or GFP fluorescent tags and pT-REx-mCherry were prepared as previously described [25, 26].

### Cell lines

Neuro-2a cells, obtained originally from the American Type Culture Collection (ATCC), were maintained in Opti-MEM (Life Technologies). The medium was supplemented with 10% v/v fetal calf serum, 1 mM glutamine, and 100 Unit mL^-1^ penicillin and 100 µg mL^-1^ streptomycin, and cells were kept in a humidified incubator with 5% v/v atmospheric CO_2_ at 37 °C.

### Transfections

Neuro2a cells were transiently transfected with the vectors using Lipofectamine 2000 reagent (Life Technologies). Specific transfection conditions for the different culture vessel types at densities of 9 × 10^4^ (Ibidi 8-well µ-chamber) or 6 × 10^6^ (T75 flasks). The following day cells (confluency of 80 – 90 %) were transiently transfected with 1.25 or 60 µL Lipofectamine 2000 and 0.5 or 24 µg vector DNA, respectively, as per the manufacturer’s instructions (Life Technologies). The next day, the medium was changed to Opti-MEM, and for the time course the medium was refreshed daily.

### Confocal imaging

Cells co-transfected with EGFPC2-GA_101_ and Httex1Q_97_-mCherry were fixed 24 h after transfection in 4% w/v paraformaldehyde for 15 min at room temperature. Nuclei were counterstained with Hoechst 33342 at 1:200 dilution (Thermo Fisher Scientific, San Jose, CA) for 30 min then washed twice in phosphate buffered saline (PBS). Fixed cells were imaged on a Leica SP5 confocal microscope using HCX PL APO CS 40× or 63× oil-immersion objective lens (NA 1.4) at room temperature. Laser used: 405 nm excitation, 445–500 nm emission– Hoechst 33342; 488 nm excitation, 520–570 nm emission–GFP; 561 nm excitation, 590 nm emission– mCherry. Single colour controls were used to establish and adjust to remove bleed through of the emission filter bandwidths. FIJI version of ImageJ [27] and Inkscape software were used for image processing.

### Longitudinal live-cell imaging

Neuro2a cells were co-transfected with pT-REx-mCherry and either EGFPC2-GA_101_ or pT-REx-Httex1Q_97_ -GFP. Medium was refreshed 24 hr post transfection and cells were imaged longitudinally with a JuLI-stage fluorescence microscope (NanoEnTek) at 15 min intervals for 96 hours. Channels used: GFP for EGFP (466/40 nm excitation, 525/50 nm emission), RFP for mCherry (525/50 nm excitation, 580 nm emission). Measurement of time of inclusion formation were extracted from files generated with automated imaging using the FIJI version of ImageJ [28]. Image processing was performed with the FIJI version of ImageJ [27] and visual inspection. Differences in inclusion formation rates were assessed by survival curve analysis in GraphPad Prism 7.05 (Graphpad Software Inc., San Diego, CA).

### Purification of PolyGA and polyQ Aggregates

Neuro-2a cells expressing either GFP-tagged 101xGA or Httex1Q_97_ in 3 replicates were harvested by pelleting (200 *g*; 5 min; 24 °C) 24 h post transfection. Cell pellets were resuspended in lysis buffer (20 mM Tris, pH 8.0; 2 mM MgCl_2_; 150 mM NaCl; 1% (w/v) Triton X-100; 20 Units/mL Benzonase, Novagen; 1× complete mini-protease cocktail; Roche) and then incubated for 30 min on ice. Lysates were diluted 2 times with PBS supplemented with protease inhibitor and aggregates were pelleted at 1000 *g* for 6 minutes. The aggregates were washed twice with 1 mL PBS, then resuspended in 1 ml PBS and subjected to fluorescence-activated cell sorting (FACS) on a BD FACS Aria III instrument with an outlet nozzle of 100 µm in diameter. The flow rate was adjusted to ∼500 events/min, and EGFP fluorescence was monitored for sorting. Sorted aggregates were pelleted (12,000 *g*; 5 min; 4 °C), resuspended in PBS and washed 3 times by pelleting as above and resuspension in PBS. The final pellets were harvested by pelleting (21,000 *g*, 6 min, 4 °C) and dissolved in 10 µL neat formic acid for 30 min at 37 °C, vortexed for 20 seconds and sonicated for 1 min three times then incubated in a shaking microfuge tube incubator (30 min, 37 °C). Samples were neutralized to pH 7.0 by titration with unbuffered 3 M Tris. The protein concentration in the sample was determined by a Bradford assay using bovine serum albumin as mass standard. A total protein of 200 µg was further processed for mass spectrometry analysis.

### Collection of cells by Pulse Shape Analysis

To assess the impact of polyGA aggregation on whole proteome, Neuro2a cells expressing GFP-tagged polyGA in 3 replicates were harvested 48 h post transfection by resuspension in PBS with a cell scraper. Cells were pelleted (120 *g*; 6 min) and resuspended in 2 mL PBS supplemented with 10 units/mL DNase I and filtered through 100 μm nylon mesh before analysis by flow cytometry. DAPI or Sytox (Thermo Fisher Scientific) was spiked into cell suspensions just before sorting to stain dead cells. Cells were analyzed by a FACS ARIA III cell sorter (BD Biosciences) equipped with 405-nm, 488-nm, 561-nm and 640-nm lasers. Live cells were gated using side and forward scatter as described previously [29]. Cells were further gated into cells with polyGA_101_ in the soluble form (ni) and those with polyGA_101_ inclusions (i) by pulse shape analysis (PulSA) as previously described [29]. Each gate recovered between 0.8-1 × 10^6^ cells which were sorted directly into PBS and then snap frozen in liquid nitrogen and stored at – 80 °C until used.

### Sample preparation for whole proteome analysis

Sorted cell populations were thawed and resuspended in 100 µl RIPA lysis buffer (25 mM Tris-HCl, pH 7.4, 150 mM NaCl, 1% v/v NP-40, 0.1% w/v SDS, 1% w/v sodium deoxycholate, 1× complete mini-protease mixture; Roche), and incubated on ice for 30 min. The concentration of proteins was measured by the Pierce microBCA Protein Assay according to the manufacturer’s instruction (Thermo Fisher Scientific). Equal amounts of protein for each sample were precipitated with six volumes of pre-chilled (−20 °C) acetone, and incubation overnight at –20°C. Samples were then pelleted (21,000 *g*, 10 min, 4 °C). Acetone was decanted without disturbing the protein pellet. The pellets were washed once with pre-chilled acetone then allowed to dry for 10 min. The protein precipitates were resuspended in 100 µl 0.1 M triethylammonium bicarbonate (TEAB) and were vortexed and then sonicated 3 times for 30 s. The samples were further processed for mass spectrometry analysis.

### Protein sample preparation for mass spectrometry

Proteins were subjected to reduction with 10 mM tris(2-carboxyethyl)phosphine (TCEP), pH 8.0, and alkylation with 55 mM iodoacetamide for 45 min, followed by trypsin digestion (0.25 µg, 37 °C, overnight). The resultant peptides were adjusted to contain 1% v/v formic acid then desalted by solid-phase extraction with an SPE cartridge (Oasis HLB 1 cc Vac Cartridge, Waters Corp., Milford, MA) pre-washed with 1 ml of 80% v/v acetonitrile (ACN) containing 0.1% v/v trifluoroacetic acid (TFA) and equilibrated with 1.2 ml of 0.1% v/v TFA three times. Samples were then loaded on the cartridge and washed with 1.5 ml of 0.1% v/v TFA before being eluted with 0.8 ml of 80% v/v ACN containing 0.1% v/v TFA and collected in 1.5 ml microcentrifuge tubes. Peptides were then lyophilized by freeze drying (Virtis, SP Scientific, Warminster, PA). The peptides were resuspended in 100 µl distilled water and quantified using microBCA assay with bovine serum albumin as the mass standard. Then, 10 µg of each sample (in a volume of 50 µl containing 100 mM TEAB) were differentially labelled by reductive dimethyl labelling using equal volumes (2 µl) of 4% light formaldehyde (CH_2_O) or 4% medium formaldehyde (CD_2_O, 98% D) and 0.6 M Sodium cyanoborohydride (NaCNBH_3_). The peptide solutions were incubated on an Eppendorf Thermomixer (Eppendorf South Pacific Pty. Ltd., Macquarie Park, NSW, Australia) at room temperature for 1 h. After quenching with 8 µl of 1% v/v ammonium hydroxide followed by further quenching with 8 µl of neat formic acid, dimethyl-labelled peptides were combined in equal amounts prior to liquid chromatography-nano electrospray ionization-tandem mass spectrometry (LC-nESI-MS/MS) analysis.

### NanoESI-LC-MS/MS analysis

Peptides were analyzed by LC-nESI-MS/MS using an Orbitrap Lumos mass spectrometer (Thermo Fisher Scientific) fitted with nanoflow reversed-phase-HPLC (Ultimate 3000 RSLC, Dionex, Thermo Fisher Scientific). The nano-LC system was equipped with an Acclaim Pepmap nano-trap column (Dionex - C18, 100 Å, 75 μm × 2 cm) and an Acclaim Pepmap RSLC analytical column (Dionex - C18, 100 Å, 75 μm × 50 cm, Thermo Fisher Scientific). For each LC-MS/MS experiment, 1 μg (whole proteome) or 0.135 μg (aggregate proteome) of the peptide mix was loaded onto the enrichment (trap) column at a flow of 5 μl/min in 3% CH_3_CN containing 0.1% v/v formic acid for 6 min before the enrichment column was switched in-line with the analytical column. The eluents used for the LC were 5% DMSO/0.1% v/v formic acid (solvent A) and 100% CH_3_CN/5% DMSO/0.1% formic acid v/v. The gradient used was 3% v/v B to 20% B for 95 min, 20% B to 40% B in 10 min, 40% B to 80% B in 5 min and maintained at 80% B for the final 5 min before equilibration for 10 min at 3% B prior to the next analysis.

The mass spectrometer was operated in positive-ionization mode with spray voltage set at 1.9 kV and source temperature at 275 °C. Lockmass of 401.92272 from DMSO was used. The mass spectrometer was operated in the data-dependent acquisition mode, with MS spectra acquired by scanning from m/z 400–1500 at 120,000 resolution with an AGC target of 5e5. For MS/MS, the “top speed” acquisition method mode (3 s cycle time) on the most intense precursor was used whereby peptide ions with charge states ≥2 were isolated with an isolation window of 1.6 m/z and fragmented with high energy collision (HCD) mode, with a stepped collision energy of 30 ± 5%. Product ion spectra were acquired in the Orbitrap at 15,000 resolution. Dynamic exclusion was activated for 30s.

### Proteomic data analysis

Raw data were analyzed using Proteome Discoverer (version 2.3; Thermo Scientific) with the Mascot search engine (Matrix Science version 2.4.1). Database searches were conducted against the Swissprot *Mus musculus* database (version 2016_07; 16794 proteins) combined with common contaminant proteins. GFP sequence (UniProt ID: P42212) was also concatenated to the Httex1Q_97_ and PolyGA_101_ sequences. Search was conducted with 20 ppm MS tolerance, 0.2 Da MS/MS tolerance and 2 missed cleavages allowed. Variable modifications were used for all experiments: oxidation (M), acetylation (Protein N-term), dimethylation (K), dimethylation (N-Term), 2H(4) dimethylation: (K) and 2H(4) dimethylation (N-term). A fixed modification used for all experiments was carbamidomethyl (C). The false discovery rate (FDR) was calculated by the Percolator node in Proteome Discoverer v 2.3.0.81 and was set to 0.5 % at the peptide identification level and 1 % at the protein identification level. Proteins were filtered for those containing at least two unique peptides in 3 biological replicates. Peptide quantitation was performed in Proteome Discoverer v.2.3 using the precursor ion quantifier node. Dimethyl labelled peptide pairs were established with a 2 ppm mass precision and a signal to noise threshold of 3. The retention time tolerance of isotope pattern multiplex was set to 0.6 min. Two single peak or missing channels were allowed for peptide identification. The protein abundance in each replicate was calculated by summation of the unique peptide abundances that were used for quantitation (light or medium derivatives). Missing quantitation values were replaced with a constant (zero-filling). The peptide group abundance and protein abundance values were normalized to account for sample loading. In brief, the total peptide abundances for each sample was calculated and the maximum sum for all files was determined. The normalization factor was the factor of the sum of the sample and the maximum sum in all files. After calculating the normalization factors, the Peptide and Protein Quantifier node normalized peptide group abundances and protein abundances by dividing abundances with the normalization factor over all samples. The normalized protein abundances were imported into Perseus software (v 1.6.5.0). Protein abundances were transformed to log2 scale. The samples were then grouped according to the replicates. For pairwise comparison of proteomes and determination of significant differences in protein abundances, paired Student’s t test based on permutation-based FDR statistics was then applied (250 permutations; FDR = 0.05; S0 = 0.1). This was justified on the basis the proteomics abundance data is normally distributed.

### Bioinformatics

Protein interaction networks were generated using Cytoscape 3.7.1[30] built-in STRING (v11.0) [31] using active interaction sources parameters on for Experiments, Databases, Co-expression neighborhood, Gene Fusion and Cooccurrence. The minimum required interaction score setting was 0.9 (highest confidence). The corresponding enriched GO annotation terms were determined by calculating their enrichment *P*-value, which we compute using a Hypergeometric test, as explained in [32]. The *P*-values are corrected for multiple testing using the method of Benjamini and Hochberg [33]. Selected GO terms were used to manually re-arrange nodes and were added to protein interaction network using Inkscape. IUPred [34] were applied to predict the intrinsically unstructured/disordered regions of proteins significantly enriched in polyGA or Httex1Q_97_ aggregates. Glutamine content was analyzed with the web-server COPid [35] (http://www.imtech.res.in/raghava/copid/). A control set of 100 random proteins (Table S1) was generated from a list of the mouse proteome obtained from the Uni-ProtKB database (http://www.uniprot.org/uniprot/?query=reviewed:yes+AND+organism:10090&random=yes). The Mann-Whitney-Wilcoxon test was employed to determine significant differences.

### Statistical Analysis

The details of the tests were reported in the figure legends. All statistical analyses were performed with GraphPad Prism v 7.05 (Graphpad Software Inc., San Diego, CA). Significant results were defined on the figures for *P* < 0.05.

### Data availability

The MS proteomic data have been deposited to the ProteomeXchange Consortium via the PRIDE [36] partner repository with the dataset identifiers PXD018505 for aggregate proteome data and PXD018824 for whole proteome data.

## RESULTS & DISCUSSION

Previously we found that polyGA_101_ as a fusion to GFP formed cytosolic inclusions in Neuro2a cells when transiently transfected. To further investigate the rate that this occurs, we used live cell imaging to track cells from 24 h after transfection onwards. Of cells with detectable levels of expression by 24 h almost all of them had formed inclusions by 60 h (Fig 1A). This was faster than comparable experiments with Httex1Q_97_ as a fusion to mCherry, which is well known to extensively aggregate in cell culture [37, 38] (Fig 1A). We noted that some lower-expressing cells showed detectable expression of polyGA only after 24 h (which we did not track) and that these were likely to form aggregates more slowly. When we co-expressed Httex1Q_97_ as a fusion to mCherry, we found the polyGA_101_ and Httex1Q_97_ formed discrete inclusions in the same cell with no apparent colocalization (Fig 1B). This suggested that any concomitant co-aggregation patterns that arise with endogenous proteins may involve distinct proteins.

**Figure 1.**
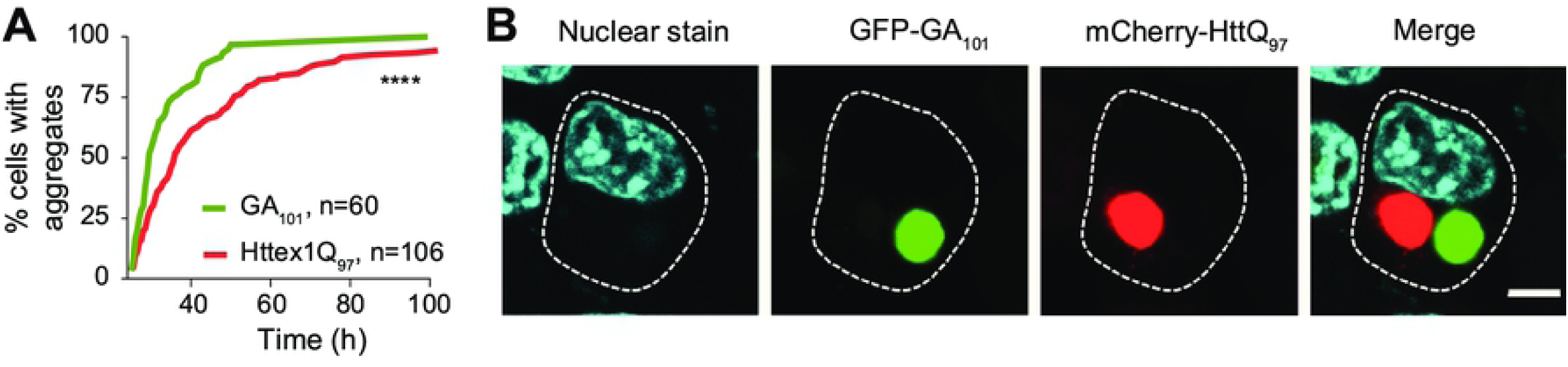
Httex1Q_97_ and polyGA_101_ rapidly form distinct inclusions in neuro2a cells. **A**. Kaplan-Meier curves of polyGA- or Httex1Q_97_ expressing cells that form visible inclusions as assessed by longitudinal imaging. Cells were tracked from 24 h post-transfection. *P*-values correspond to log-rank (Mantel-Cox) test. **B**. Confocal micrographs of neuro2a cells co-expressing GFP-tagged GA_101_ and mCherry-tagged Httex1Q_97_, fixed 24 hr post-transfection and stained with Hoechst33258 (cyan) to visualize nuclei. Scale bar represents 5 µm.

To investigate these potential differences, pellets recovered from lysates of neuro2a cells expressing GFP-tagged Httex1Q_97_ or GFP-tagged polyGA_101_ were sorted to purify the aggregates using flow cytometry via monitoring the GFP fluorescence. Quantitative proteomics, by way of dimethyl isotope labelling, was used to define the proteins enriched in each aggregate class after normalization to total mass of protein. We observed 737 proteins. Of these 63 were significantly enriched in polyGA inclusions (3 replicates, a permutation-based FDR cut-off of 5% and S0 of 0.1) and 48 were enriched in Httex1Q_97_ (Table 1, Table S2 and Fig 2A).

**Table 1:**
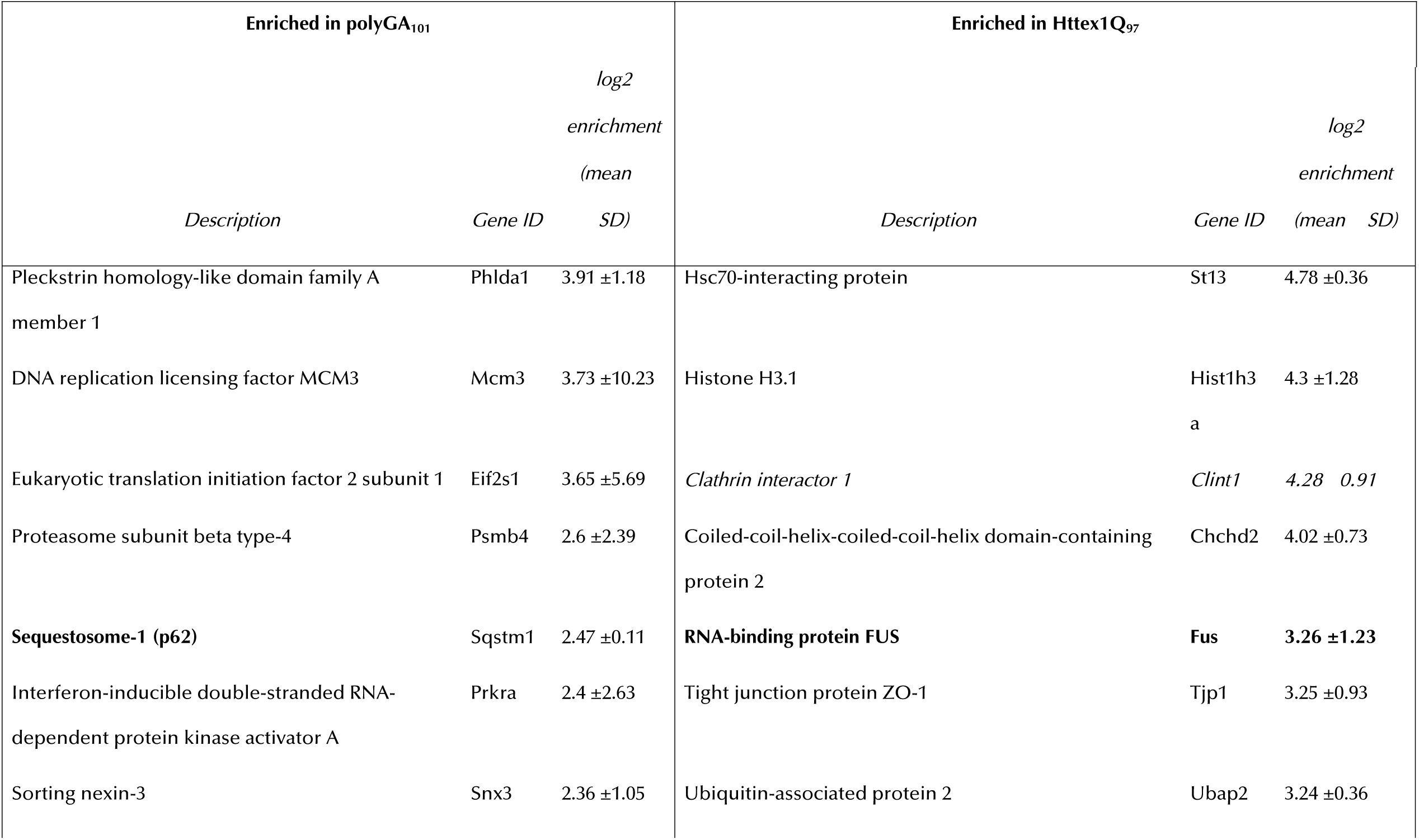
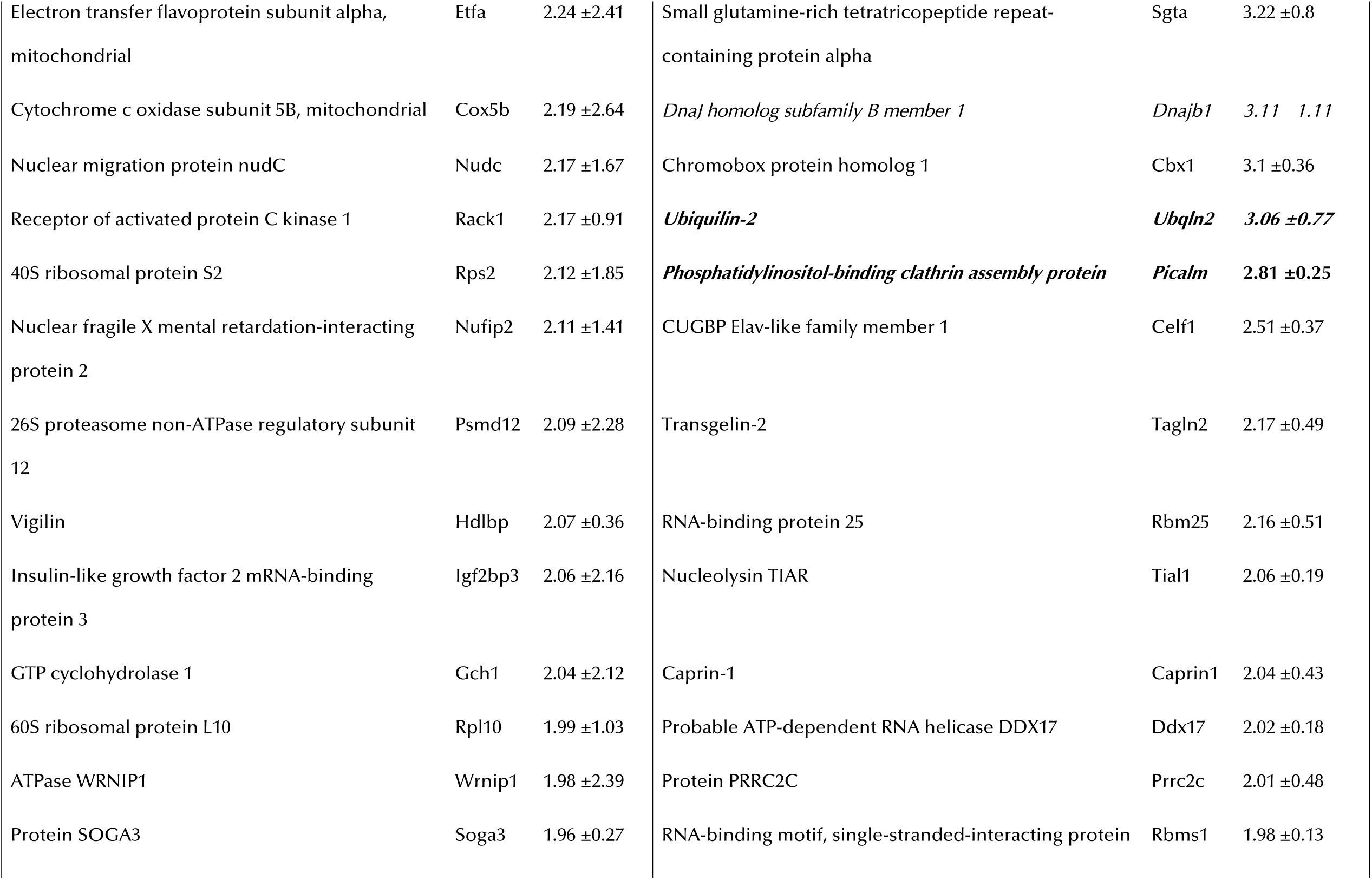
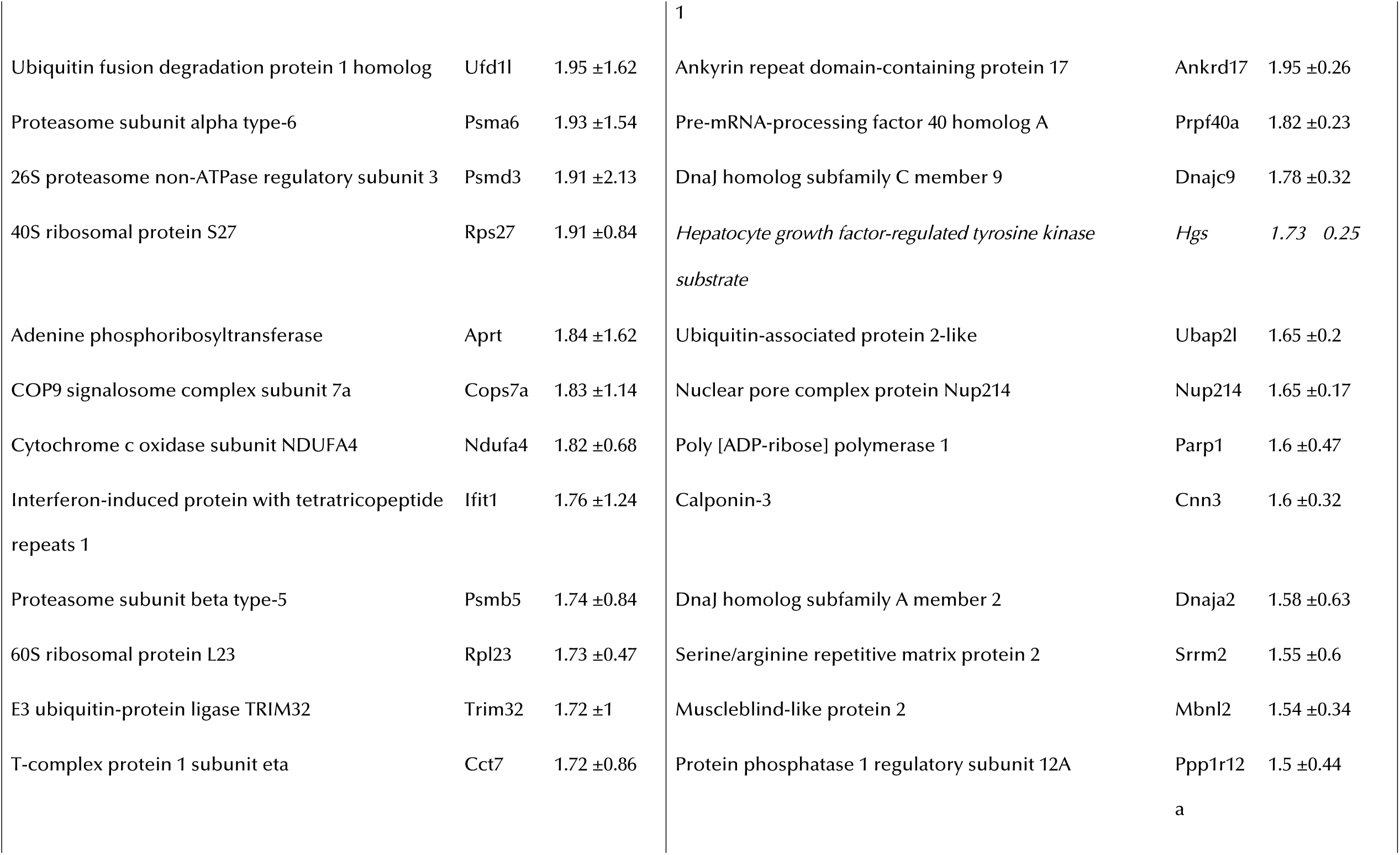
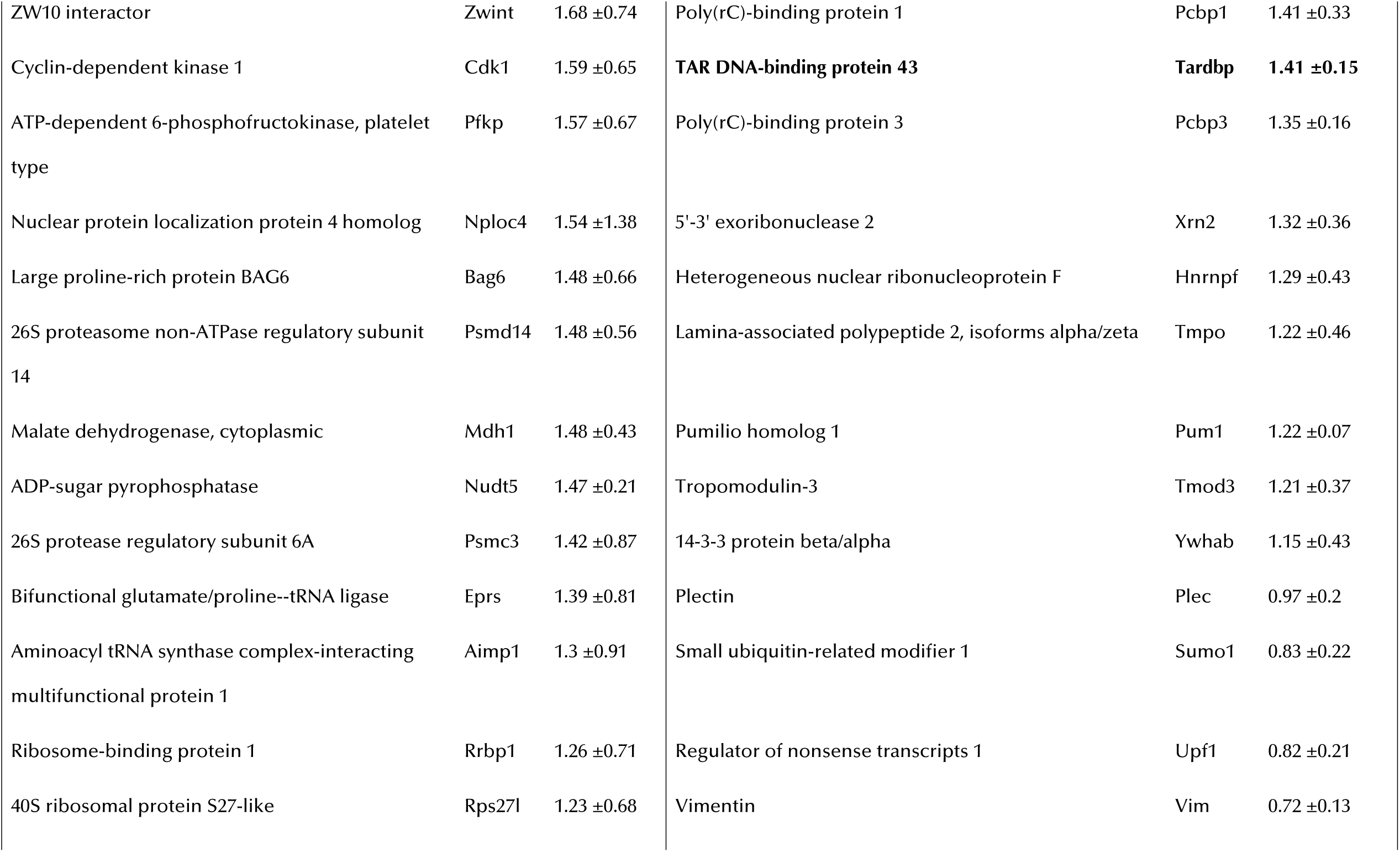
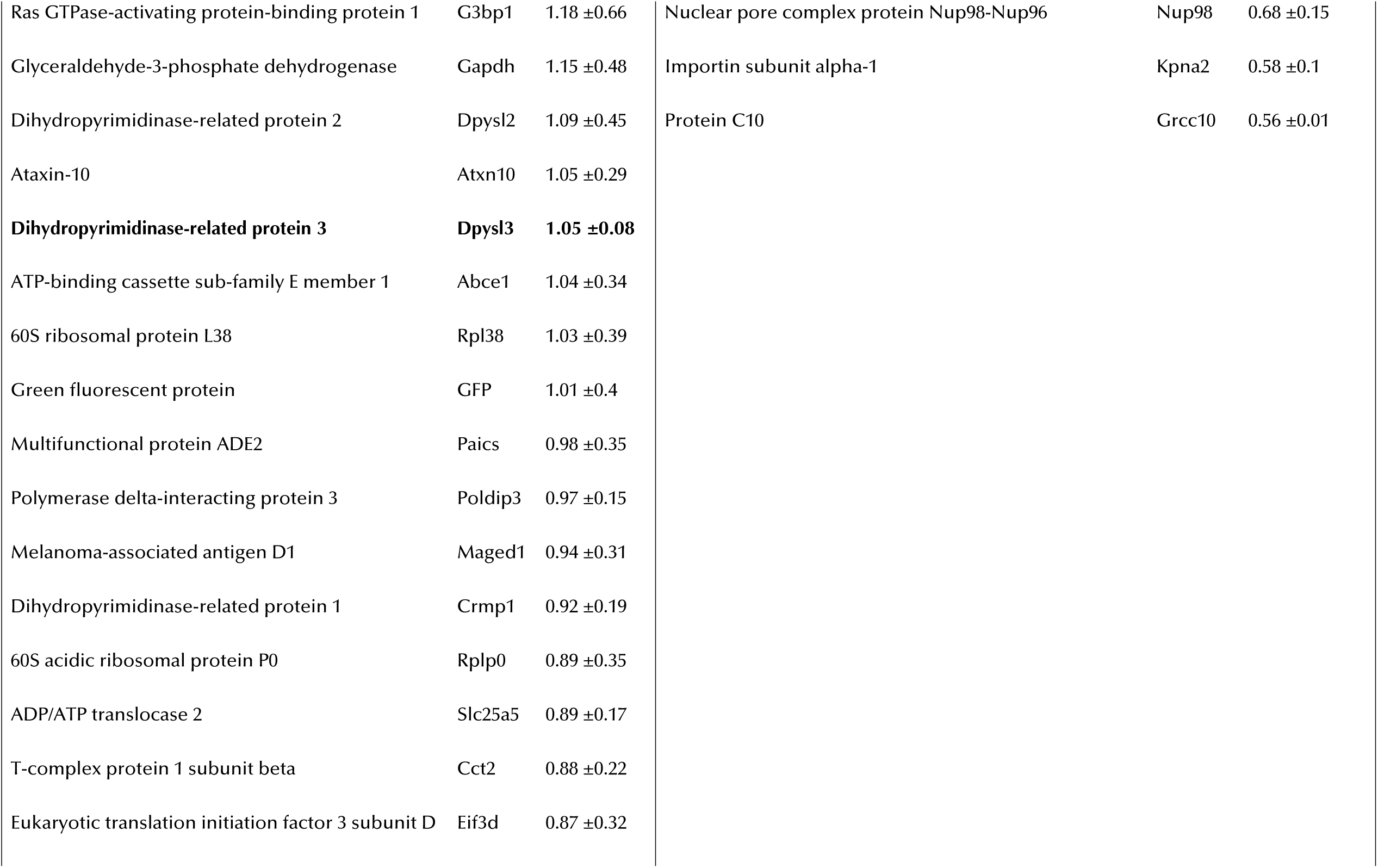
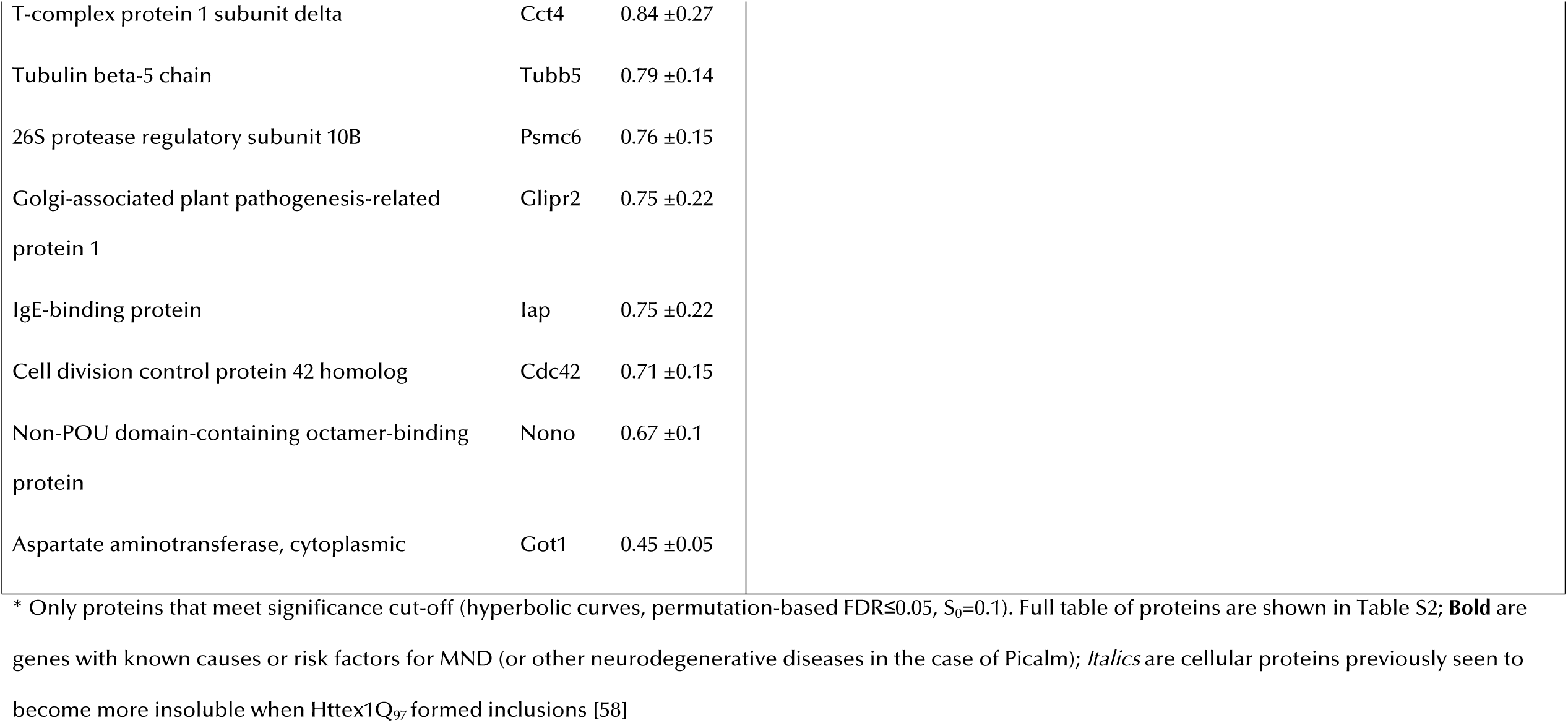
Proteins enriched in inclusions of polyGA_101_ and Httex1Q_97_*

**Figure 2.**
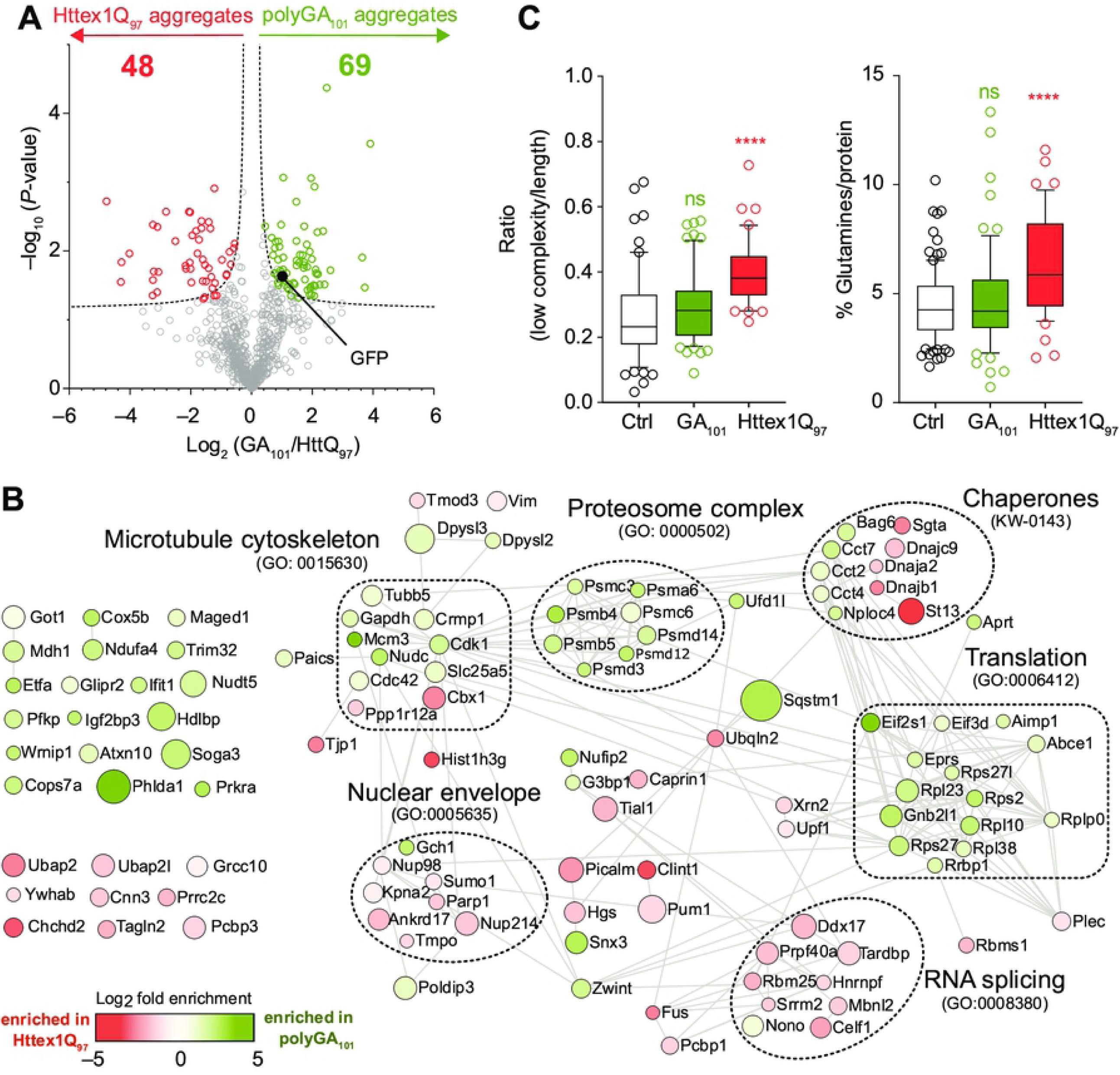
Proteome recruitment patterns to polyGA_101_ and Httex1Q_97_ inclusions. **A**. Volcano plot of proteins identified in the inclusions. *P*-values were calculated by a two-sided one samples t-test with null hypothesis that abundances were unchanged and the Log_2_ ratio was equal to 0. Proteins meeting stringency thresholds (hyperbolic curves, FDR≤0.05, S_0_=0.1) are shown as colored circles. **B**. STRING interaction maps (v.11) determined in Cytoscape (v3.7) for proteins significantly enriched in the inclusions (the full list of proteins are in Table S2). The analysis was done at the highest confidence setting. Each protein was represented by a colored circle sized proportionally to −log_10_ (*P*-value). The color scale represents logarithm of fold change. Selected significantly enriched GO terms (GOCC, GOPB, and UniProt keywords) are displayed (Full terms are shown in Table S3). **C**. Analysis of enriched proteomes for low-complexity regions (IUPred-L) and high glutamine content. Significance of difference was assessed against a control dataset of random mouse proteins (Table S1) with the Mann–Whitney–Wilcoxon test. Whiskers extend from 10 to 90%.

We observed notable features in this dataset consistent with known pathological markers of polyGA inclusions. Namely in C9ORF72 mediated MND, a subset of inclusions is non-reactive to TDP-43 [39]. In most other forms of MND, TDP-43-reactive inclusions are a key pathological signature of neurons in disease [40]. These TDP-43 negative inclusions were previously found to be immunoreactive for polyGA, suggesting they form by polyGA aggregation [41, 42]. We observed a lack of TDP43 in the polyGA inclusions by virtue of an enrichment in the Httex1Q_97_ inclusions (Table 1). In addition, the TDP43-negative inclusions seen in vivo are immunoreactive to p62 [43] and lack immunoreactivity to FUS, optineurin, alpha-internexin and neurofilament [44, 45]. In our data p62 (also called sequestesome 1) is one of the most enriched proteins in polyGA inclusions, whereas Fus appeared excluded by virtue of its enrichment in Httex1Q_97_ inclusions, which has been observed previously in cell models of polyQ aggregation and human pathology [25, 46-48]. Hence these data point to the cell model of polyGA inclusions mimicking the process of aggregation and recruitment seen in vivo and also providing specificity of co-recruitment relative to Httex1Q_97_.

Analysis of the differences is shown visually in Fig 2B by a (STRING) protein-protein interaction map and annotation to functional networks. Overall both inclusions yielded an enrichment for gene ontology and KEGG networks of microtubule cytoskeleton, proteasome complex, chaperones, RNA splicing and nuclear envelope (Fig 2C; Table S3). These findings are in accordance with prior findings that protein aggregation impacts these biological processes and in particular an involvement in machinery for their clearance and degradation [49-52].

In addition, the data points more directly to proteins and genes implicated in MND phenotype and mechanisms. Phlda1 was one of the proteins enriched in polyGA inclusions. Previously it was found that Phlda1 was upregulated in Fus-mutant motor neurons and that this was an adaptive response to protect against apoptosis [53]. Phlda1 was also observed upregulated in sporadic MND fibroblasts treated to stress compared to controls [54]. Nudt5 was also found enriched in inclusions, and expression of this gene was significantly increased in motor neurons derived from induced plutripotent cells from MND patients over controls [55]. Another protein of note enriched in the polyGA aggregates was Dpysl3. A missense mutation that has been linked to MND risk in French population and in culture expression of the mutation leads to shorted neuronal survival [56]. Hence it remains plausible that co-aggregation of these proteins into polyGA inclusions sequesters their activity and renders cells less resilient to stress triggers.

The Httex1Q_97_-enriched proteome also yielded noteworthy findings. Previously it was found that polyQ can preferentially co-recruit proteins containing intrinsically disordered domains and proteins enriched in glutamine (IDRs) [25, 47]. These patterns were also observed in our data (Fig 2C). However, polyGA did not show these enrichment patterns, indicative of specificity for polyQ in recruiting IDRs and Q-rich proteins.

To assess whether the changes in polyGA inclusion formation had other effects on proteome abundance, we expressed polyGA_101_ and at 48 h after transfection sorted live cells into those with visible aggregates from those without by flow cytometry sorting method and pulse shape analysis [57] (Fig 3A). We found cells with inclusions were more reactive to Sytox (Fig 3A inset), which is indicative of dying and dead cells, than cells without inclusions so we excluded these cells from analysis. 35% of the remaining live cells expressing polyGA had inclusions (Fig 3A). This was a lower yield of aggregates than we measured by live cell imaging at 48h in Fig 1A (about 90%), which we attribute to this experiment being more inclusive of the lower-expressing cells and other differences in the experimental conditions that affect aggregation rates such as phototoxicity of live cell imaging. Nonetheless, the yield obtained made the experiment amenable for comparing cells with versus without inclusions. Out of 2420 proteins identified, we observed 56 proteins that significantly changed abundance in these sorted cell populations (Fig 3B; Table S4). There was no overlap in the proteins seen enriched in polyGA inclusions with proteins that changed expression due to polyGA aggregation. This provides firmer confidence that the enrichment seen in the polyGA aggregates arises from co-aggregation rather than changes in gene expression. Of the genes that changed expression, protein interaction networks yielded significant enrichment in networks including nuclear speck (GO: 0016607), ribosome biogenesis (GO: 0042254), chromosome (GO:0005694), mitochondrion (GO:0005739) and Golgi-to-ER-traffic (MMU-6811442) (Table S5). These pathways would be anticipated to be activated by stress responses incurred by protein aggregation, however, we did not note any striking changes that pertained to novel mechanisms other than that from this data.

**Figure 3.**
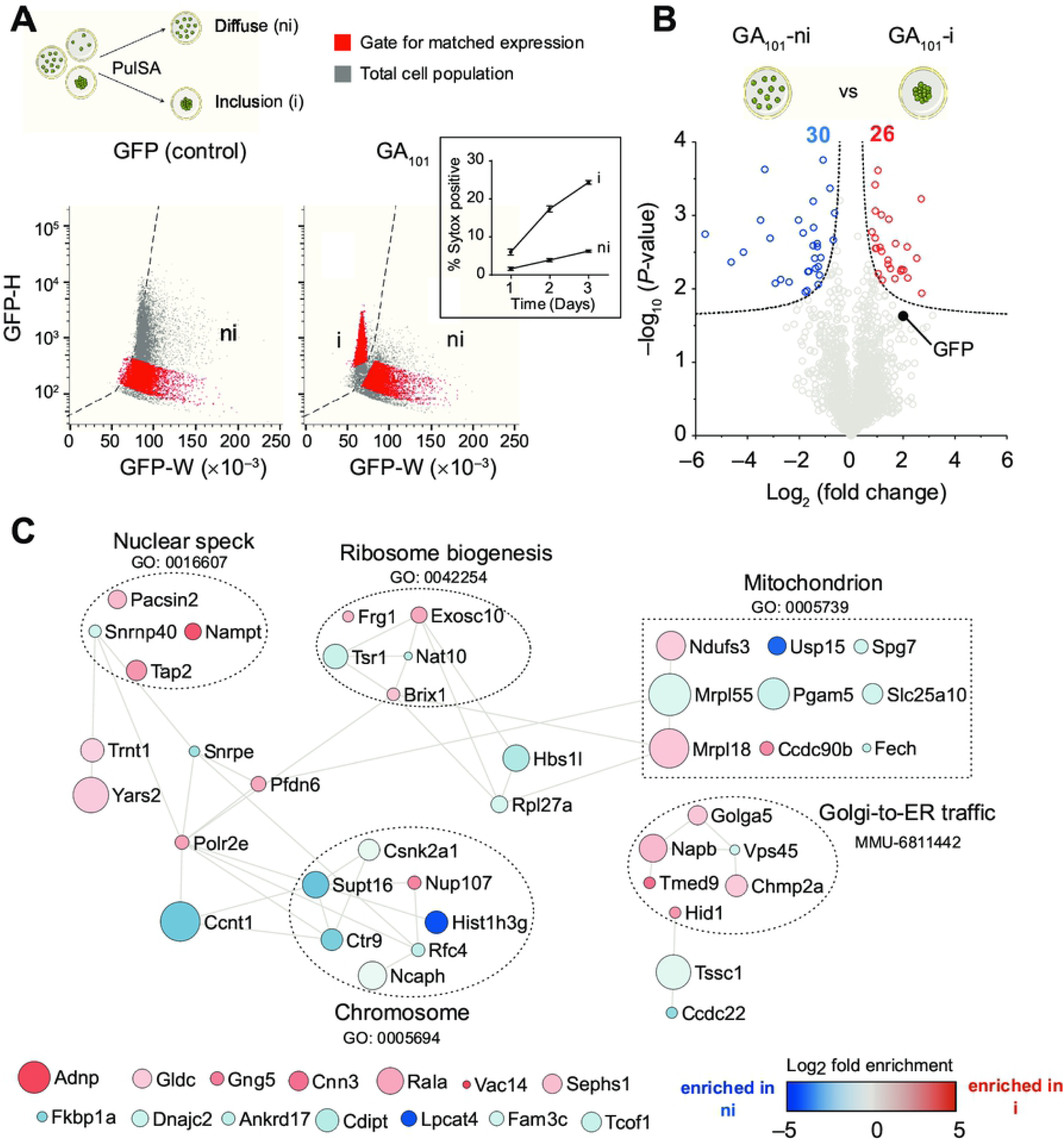
Cellular protein abundance changes arising from polyGA_101_ aggregation. **A**. Schematic of flow cytometry method of pulse shape analysis (PulSA) to sort cells enriched with inclusions (i) from those without inclusions (ni). Cells with inclusions display shorter width (W) fluorescence values versus cells with soluble protein, and typically higher height values (H) arising from the condense foci of fluorescence inside the cells. Cells were sorted to exclude dead cells by Sytox reactivity. Inset shows percentage of transfected cells reactive to Sytox by time after transfection. *n*=4, means ± SD shown. **B**. Volcano plots of proteins that changed their abundance upon polyGA aggregation. The data for all proteins are plotted as log2-fold change versus the −log10 of the *P*-value. The dotted line indicates significance cut-off (hyperbolic curves, FDR≤0.05, S_0_=0.1) and proteins meeting stringency thresholds are shown as colored circles. **C**. Protein-protein interaction network (STRING v11) of proteins significantly changed in abundance upon polyGA aggregation (the full list of proteins are in Table S4). The analysis was done at the highest confidence setting. Each protein was represented by a colored circle sized proportionally to −log_10_ (*P*-value). The color scale represents logarithm of fold change. Selected significantly enriched GO terms (GOCC, GOPB, and UniProt keywords) are displayed (Table S5).

Lastly we investigated the overlap of proteins enriched in Httex1Q_97_ inclusions with our previously reported changes in solubility of whole cell proteome before versus after inclusions had formed [58]. In that dataset we observed 17 proteins that significantly decreased in solubility as cells expressing Httex1Q_97_ shifted from a dispersed unaggregated state to forming inclusions [58]. Of these, 5 proteins were found in our list of 48 proteins significantly enriched in Httex1Q97 inclusions (Picalm, Hgs, Clint1, Ubqln2 and Dnajb1). Four of these proteins (all except Dnajb1) form a robust protein-protein interaction network with a significant gene ontology enrichment for clathrin coat assembly (GO:0048268; FDR of 0.0031) suggestive that this mechanism is involved in polyQ aggregation. Clint1 and Ubqln2 were previously shown to colocalize to polyQ inclusions, supporting this conclusion [47, 59]. An interesting note with respect to mechanism is that UBQLN2 targets ubiquitinated substrates for degradation in ERAD and autophagy [60]. Furthermore mutations in UBQLN2 cause MND, and appear to operate by impairing protein degradation of ubiquitinated proteins [61]. Further supporting an important role linking protein aggregation, degradation more broadly to these neurodegenerative diseases is the enrichment of Picalm in the polyQ inclusions. Picalm is an phosphatidylinositol-binding clathrin assembly protein and has been shown via GWAS as a top ten risk for Alzheimer’s disease [62, 63]. It has been reported to modulate intracellular APP processing and plaque pathogenesis [64], modulate autophagy and alter tau clearance [65].

Collectively the data here reports proteins that co-aggregate into two very different neurodegenerative disease proteinaceous deposits. The findings provide specificity of proteins to the aggregation type that provide useful perspective to that reported by others. Moreover, the mechanisms of protein clearance mechanism appear relevant to both aggregation types and notably of a number of proteins in the Httex1Q_97_ aggregates that when mutated are modifiers of MND risk. Therefore, the findings identify a synergy of biological mechanisms involved in protein degradation that appear central to at least two different neurodegenerative diseases, and possibly more applicable to the other neurodegenerative diseases involving inappropriate protein aggregation.

## ACKNOWLEDGMENTS

We thank the Bio21 Melbourne Mass Spectrometry and Proteomics facility. This work was funded by grants to DMH (National Health and Medical Research Council APP1161803 and Motor Neuron Disease Research Institute, Australia small grant) and to DMH and GER (Australian Research Council DP170103093). MR acknowledges support from an Australian Government Research Training Program (RTP) Scholarship and an Egyptian Ministry of Higher Education and Scientific Research PhD scholarship.

## SUPPORTING INFORMATION CAPTIONS

**Table S1. List of random proteins from mouse Uniprot database**.

**Table S2. Proteins enriched in inclusions of polyGA**_**101**_ **and Httex1Q**_**97**_. **Relates to Table 1 & Fig 2**.

**Table S3. Gene Ontology terms enriched among proteins identified in polyGA**_**101**_ **or Httex1Q**_**97**_ **inclusions. Relates to Fig 2**.

**Table S4. Cellular abundances of proteins caused by polyGA**_**101**_ **aggregation. Relates to Fig 3**.

**Table S5. Gene Ontology terms enriched among proteins that changed abundance upon polyGA**_**101**_ **aggregation. Relates to Fig 3**.

